# PDH Mediated Mitochondrial Respiration Controls the Speed of Muscle Stem Cell Activation in Muscle Repair and Aging

**DOI:** 10.1101/2020.02.14.950162

**Authors:** Manmeet H. Raval, Pin-Chung Cheng, Nicholas Guardino, Sanjana Ahsan, Hao Zhou, Rajiv Lochan Tiwari, Lu Wang, Andrew Chareunsouk, Maxwell Ederer, Ara B. Hwang, Matt Ellenberger, Rob Pepin, Daniel Raftery, Daniel Promislow, Keyue Shen, Andrew S. Brack, Joseph T. Rodgers

## Abstract

Decline in the skeletal muscle stem cell (MuSC) function is a major contributor to age-associated impairments in muscle regeneration and function. The ability of MuSCs to activate (i.e. exit quiescence, enter the cell cycle, and divide) following injury is a critical step that initiates muscle regeneration. However, the mechanisms that regulate MuSC activation function are poorly understood. Here, we show that the activation function, specifically the speed by which cells progress through G0-G1, declines tremendously with age in mouse MuSCs. Using a number of *in vivo* models and *ex vivo* assays of MuSC activation and muscle regenerative functions, live cell metabolic flux analyses, and metabolomics we present data indicating that changes in MuSC mitochondrial flux underlie age-associated changes in MuSC activation. We show that, in the course of MuSC activation, there is a profound,16-fold, increase in ATP production rates, which is fueled largely by increases in pyruvate flux into mitochondria. We found that MuSCs from aged mice display progressive defects in the ability to increase mitochondrial flux during activation and that this correlates with higher levels of phosphorylated, inactivated, pyruvate dehydrogenase (PDH). Importantly, we demonstrate that pharmacologic and physiologic methods to induce dephosphorylation and activation of PDH in MuSCs are sufficient to rescue the activation and muscle regenerative functions of MuSCs in aged mice. Collectively the data presented show that MuSC mitochondrial function is a central regulator of MuSC activation and muscle regenerative functions. Moreover, our results suggest that approaches to increase MuSC pyruvate oxidation may have therapeutic potential to promote muscle repair and regeneration.

## INTRODUCTION

Skeletal muscle stem cells (MuSCs), also known as satellite cells, are required for skeletal muscle repair and regeneration ^1–3^. In normal conditions MuSCs reside in muscle in a mitotically quiescent state ^4^. An injury to their host muscle induces them to activate; to exit quiescence, enter the cell cycle, and divide. The progeny of activated MuSCs subsequently proliferate and differentiate to ultimately repair or form new muscle fibers ^5,6^. The ability of MuSCs to activate and transition from a quiescent to a proliferative state is crucial for maintaining tissue repair and homeostasis ^3^. However, the regulatory mechanism of the MuSC activation step is poorly understood.

A number of previous reports have shown that the activation function of MuSCs, (i.e., the speed by which they transition from quiescence into and through the cell cycle) is regulated by a variety of cell-intrinsic and cell-extrinsic factors ^7^. High-Pax7 expressing MuSCs are slow to activate in response to injury compared to low-Pax7 expressing MuSCs ^8^. Mx1^+^ MuSCs are enriched for Pax3, have low level of reactive oxygen species (ROS) and confer stress tolerance via induction of mTORC1 ^9,10^. We and others have previously described a HGF-mTORC1 ^11,12^ and HMGB1-CXCR4 ^13^ regulated transition of stem cells into a primed, or poised, state of quiescence termed G_Alert_. The key functional distinction of stem cells in G_Alert_ is the increase in the speed of their injury-induced activation compared to nonprimed, G_0_, stem cells. Importantly, previous work also shows that there is a strong correlation between an increase in activation and muscle regenerative functions of G_Alert_ MuSCs ^11–13^.

Aging leads to a progressive loss of healing capacity which can be attributable to the loss of stem cell function ^14,15^. In aged mice, decline in MuSC function has been shown to be a major factor contributing to poor muscle repair ^16–18^. Aging also leads to major changes in MuSC mitochondrial function, metabolism, oxidative state, and epigenetic modifications ^19–21^. Recently, there have been a number of reports that have begun to dissect the impact of cellular metabolism on MuSC function. Overall, numerous studies have identified a positive correlation between MuSC oxidative capacity and several aspects of MuSC function including cell number and muscle regenerative activity ^22–24^. Several recent reports have delineated mechanisms that connect key tricarboxylic acid (TCA) cycle intermediates with transcriptional and epigenetic changes important for the function of MuSCs in muscle regeneration ^25–29^. Indeed, cellular metabolism has been linked to quiescence, self-renewal, and differentiation function in other tissue-specific stem cell systems as well ^30–33^. This clearly suggests that cellular metabolism is a powerful driver of stem cell function and that quiescent and cycling MuSCs, and many other stem cell pools, display age-associated qualitative and quantitative changes in metabolism. However, the interaction between age-associated and activation-induced changes in MuSC metabolism is only beginning to be understood.

Here, we show that MuSCs display a profound increase in mitochondrial metabolic activity during the transition from a quiescent state into the cell cycle (i.e. activation). We show that PDH regulates the speed of MuSC activation via controlling flux into the mitochondria. Moreover, we demonstrate that reduced mitochondrial flux directly contributes to age-associated decreases in MuSC activation speed and that increasing mitochondrial flux, by activating PDH or suppressing Lactate dehydrogenase-A (LDHA) activity, can improve activation speed and muscle regeneration in aged mice.

## RESULTS

### Age-associated delays in MuSC G0-G1 progression

Consistent with previous reports ^34^, we found that with increasing age, animals have a reduced ability to repair muscle damage. Seven and 14 days after intramuscular injection with BaCl_2_, a means to induce muscle injury, we found an inverse correlation between animal age and the cross-sectional area (CSA) of nascent, centrally nucleated, muscle fibers. Juvenile (1.5 - 2 months old) mice displayed a very rapid increase in fiber CSA after injury and had significantly larger fibers than adult (5 – 7 months old) and aged (22 – 24 months old) animals (Figures 1A-C). Aged animals had the smallest nascent fibers at 7 and 14 days post injury (DPI) (Fig. 1B-C).

**Figure 1:**
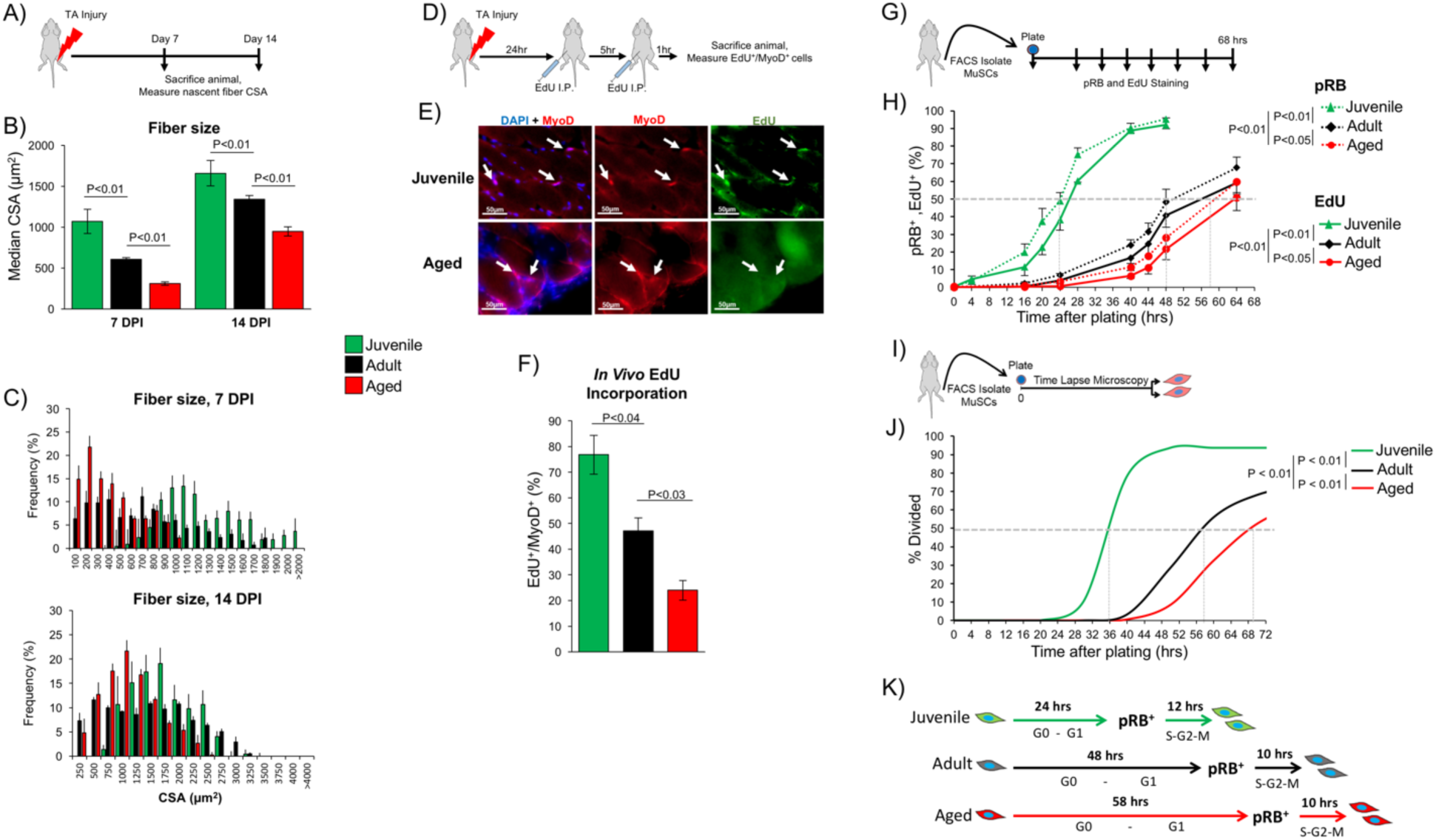
The speed of muscle stem cell activation and muscle regeneration decreases with age. A) Schematic representation of the BaCl_2_ TA injury procedure for tissue regeneration assay. B) Median fiber CSA of injured juvenile, adult, and aged mice at 7 DPI and 14 DPI. Each bar represents mean of the median CSA of centrally nucleated fibers quantified by morphometric analysis of TA cryosections ± SEM (N = 3-5). Significance was calculated by Students t-test. C) Histograms displaying the distribution of fiber CSA of the data that are summarized in Figure 1B. Each bar represents the mean ± SEM of the percentage of centrally nucleated fibers that fall within the bin range (N = 3-5). D) Schematic representation of the hemostat TA injury experimental procedure for *in vivo* EdU incorporation experiments. E) Representative IF-IHC staining of TA sections from injured mice of different age groups at 30 hours-post-injury. DAPI (blue), MyoD (red) and EdU (green). F) Quantification of EdU^+^ MyoD^+^ MuSCs in TA cryosections from injured mice of different age groups at 30 hours post-injury. Data are presented as mean ± SEM (N = 4) and significance was calculated by Student’s t-test. G) Schematic representation of *ex vivo* activation experiments analyzing pRB and EdU incorporation. H) Quantification of the percentage of MuSCs that are pRB^+^ (dotted line) and EdU^+^ (solid line) at different time-points post isolation through the course of activation. Data are presented as mean ± SEM (each data point; n = 3-7). Data from MuSCs from juvenile, adult, and aged are presented in green, black and red line, respectively. P values indicate statistical significance between age groups by 2-way ANOVA. I) Schematic representation of the *ex vivo* time to division analyses. J) Quantification of time required for first division (activation) *ex vivo*. Data represented as cumulative percentage of the MuSCs undergoing first division up to 72 hours post-isolation. MuSCs isolated from juvenile mice (green, N = 143 MuSCs), MuSCs isolated from adult mice (black, N = 91 MuSCs), and MuSCs isolated form aged mice (red, N = 191 MuSCs). P values indicate statistical significance between age groups by 1-way ANOVA. K) Model summarizing *ex vivo* activation of MuSCs from animals of different ages. Time values over the G0-G1 arrows indicate the time required for half of the MuSC population to become pRB^+^, as estimated from Figures 1H, dotted gray line. Time values over the S-G2-M arrows indicate the difference between the time required for half of the population to complete cell division, as estimated from Figure 1J, dotted gray line, and the time to become pRB^+^. MuSCs from aged mice require much more time to become pRB^+^ then MuSCs from adult or juvenile mice. However, the time from when a half a MuSC population is pRB^+^ until they complete cell division, 10-12 hours, remains consistent through the course of aging.

It has been shown that with age, MuSCs display defects and delays in their ability to activate in response to muscle injury ^7,21^. Indeed, we observed age-associated defects in the ability of MuSCs to enter the cell cycle following injury. Using incorporation of EdU nucleotide as a marker of cells that have entered S-phase of the cell cycle, we found an age-associated reduction in the fraction of MyoD^+^ MuSCs that had incorporated EdU in crush-induced injured muscles 30 hours after injury (Fig. 1D-F). In juvenile mice, we found that the 77% of activated MuSCs in the injured TA muscle were EdU^+^ while only 47% of the MyoD^+^ cells in the injured muscles of adult mice had incorporated EdU (Fig. 1F). MuSCs from aged mice showed the lowest proportion of EdU incorporation only 24% of activated MuSCs had incorporated EdU at 30 hours after muscle injury (Fig. 1F). These results suggest that with age, MuSCs develop defects in their ability to enter the cell cycle. Moreover, these results demonstrate a correlation between the ability of MuSCs to rapidly incorporate EdU after injury and the progress of the muscle regenerative response.

To gain a more detailed understanding of dynamics of MuSC activation, we utilized an *ex vivo* model of MuSC activation (Fig. 1G). Previous work has shown that FACS-mediated isolation (Fig. S1A) and purification of MuSCs from noninjured muscle closely recapitulates many aspects of *in vivo* injury-induced MuSC activation ^11,35,36^. Consistent with the *in vivo* experiments, we found that there was a delay in the progression of MuSCs through restriction point, as identified by phospho-Ser 807/811 RB (pRB)^+^ MuSCs, and entry into S-phase, as identified by EdU^+^ MuSCs (Fig. S1B). We found that half of the population of MuSCs isolated from juvenile mice were pRB^+^ after 24 hours in culture and were EdU^+^ after 27 hours (Fig. 1H). MuSCs from adult mice required more time, half of the population was pRB^+^ at 48 hours and EdU^+^ at 54 hours MuSCs from aged mice required the longest amount of time: they required 58 and 63 hours before half of the population was pRB^+^ and EdU^+^, respectively (Fig. 1H). These results show that with age, MuSCs require more time to progress through G0-G1 phases of the cell cycle and enter S-phase.

To further analyze the kinetics of MuSC activation, we used time-lapse microscopy to measure the time required by MuSCs to complete cytokinesis and divide after isolation and plating (Fig. 1I). MuSCs from juvenile mice required very little time to divide after isolation. Half of the MuSCs had divided within the first 36 hours after isolation (Fig. 1J). Consistent with the pRB and EdU analysis, we found that with increasing age, MuSCs required more time to complete the first division following isolation. The time required for half of the population of MuSCs from adult mice to divide was 58 hours, and, again, MuSCs from aged mice required the most time, 68 hours for half of the population to divide (Fig. 1J).

When we combined these data on MuSC cell cycle entry and division, we noticed a consistent pattern across all of the age groups where the time point at which half of the MuSC population became pRB^+^ occurred approximately 10-12 hours prior cell division (Fig. 1K). Because RB phosphorylation occurs at the end of G1, triggering entry into S-phase, these data suggest that the time required to complete the phases of the cell cycle following RB phosphorylation, S, G2, and M phases, is not dramatically altered in MuSCs with age. Interestingly, the phases prior to RB phosphorylation, G0-G1 phases, display a dramatic age-associated lengthening, from ∼24 hours in MuSCs from juvenile mice to ∼58 hours in MuSCs from aged mice (Fig. 1H). These results suggest that delays in G0-G1 of the cell cycle underlie age-associated delays in MuSC activation.

### Mitochondrial activity of freshly isolated MuSCs decreases with age

It is well established that cellular energy levels and metabolism regulate progression through G1 phases of the cell cycle ^37,38^. Consistent with previous work ^24^, we observed a trend of decreased MitoTracker Deep Red (MTDR) staining of MuSC from animals of increasing age, suggesting a decrease in mitochondrial function with age (Fig. 2A-B). As MTDR staining is not always a reliable reporter of mitochondrial activity in stem cells ^39^, we performed fluorescence lifetime imaging microscopy (FLIM) to more directly assess the metabolic state of freshly isolated MuSCs. This method relies on the distinct dynamics of the intrinsic fluorescence lifetime of free and bound NADH inside cells, where a greater proportion of bound NADH correlates with higher levels of flux through the mitochondrial respiratory chain ^40,41^. We found that MuSCs from juvenile mice exhibited a higher mitochondrial bound/total NADH ratio compared to adult and aged MuSCs (Fig. 2C-D). Similarly, MuSCs from adult mice exhibited significantly greater mitochondrial bound/total NADH ratios than MuSCs from aged mice. These results suggest there is an age-associated decrease in the mitochondrial function of freshly isolated MuSCs.

**Figure 2:**
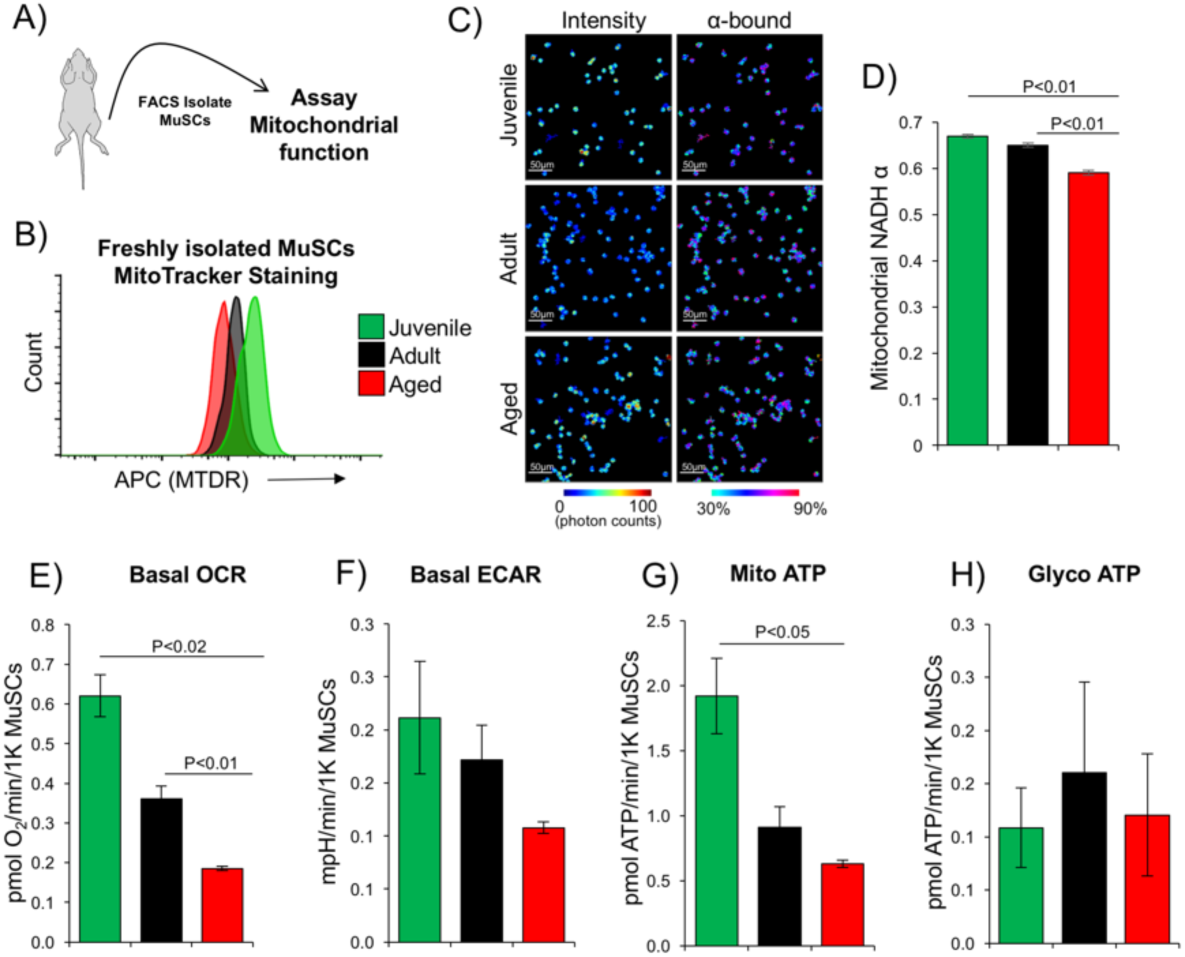
Age-associated decline in mitochondrial activity of freshly sorted MuSCs. A) Schematic representation of the experimental procedure analyzing mitochondrial function from freshly isolated MuSCs. B) Representative FACS histogram of freshly isolated MuSCs from juvenile (green), adult (black), and old (red) mice and stained with MTDR (N = 4). C) Pesudocolor-redered FLIM autofluorescent images of freshly isolated MuSCs from mice of different age groups. Color coding from the phasor histograms demonstrates NAD(P)H intensity and mitochondrial αbound assessed using phasor analysis of fluorescent lifetimes for NAD(P)H and FAD signals. D) Quantification of the mitochondrial αbound (ratio of enzyme-bound NAD(P)H over total NAD(P)H) in mitochondria. Data presented as mean ± SEM (juvenile, N = 113 MuSCs; adult, N = 139 MuSCs; aged, N = 195 MuSCs) and significance was calculated by Student’s t-test. E) Seahorse extracellular flux analyses of OCR of freshly isolated MuSCs from juvenile, adult, and aged mice. F) Seahorse extracellular flux analyses of ECAR presented in mpH/min/1K. G) Mito ATP production rates as calculated from OCR and ECAR measurements. H) Glyco ATP production rates as calculated from OCR and ECAR measurements. E-H) Data presented as mean ± SEM (N = 3) and significance was calculated by Student’s t-test.

To more directly evaluate the metabolic properties of MuSCs, we performed live cell flux analyses using a Seahorse Bioanalyzer. Consistent with the MTDR and FLIM measurements, we observed a dramatic age-associated decrease in the basal oxygen consumption rates (OCR) of freshly isolated MuSCs (Fig. 2E). MuSCs from juvenile mice displayed 2- and 3-fold higher OCR, on a per-cell basis, compared to MuSCs from adult and aged mice respectively. We also observed a similar, but lower in magnitude, trend in basal extracellular acidification rates (ECAR) (Fig. 2F). Because both LDHA, via production of lactate and H^+^, and mitochondrial respiration, via CO_2_-mediated medium acidification, are the major contributors to ECAR, we performed a series of calculations to correct for this and impute ATP production rates from the mitochondria (mito ATP) and glycolysis (glyco ATP) ^42,43^. Interestingly, we found that mito ATP production rates decreased dramatically with age (Fig. 2G) however, we observed no age-associated changes in glyco ATP production rates (Fig. 2H). Combined, these results show that mitochondrial function of freshly isolated MuSCs decreases with age.

### Aged MuSCs have a reduced ability to increase mitochondrial function upon activation

Recent studies have demonstrated that the act of FACS isolating MuSCs induces their activation and that freshly isolated MuSCs are in the early stages of activation ^44,45^. To determine if the age-associated decrease in mitochondrial function of freshly isolated MuSCs persisted, we further mapped the mitochondrial activity and respiration of MuSCs during the course of activation. Using the *ex vivo* culture model (Fig. 3A), we observed that from time of isolation to 48 hours in the course of activation, MuSCs from all age groups showed a dramatic increase in basal OCR (Fig. 3B; S2A-C). MuSCs from juvenile mice displayed both the highest respiration rates as well as the most rapid induction in mitochondrial activity compared to MuSCs from adult and aged mice (Fig. 3B). MuSCs from adult mice displayed significantly higher basal OCR than MuSCs from aged mice. We observed the same pattern of increase in MTDR staining (Fig. 3C; S2D-F). While we also observed a significant increase in basal ECAR during activation, we did not observe any clear differences between age groups (Fig. 3D). Using the OCR and ECAR measurements to calculate mitochondrial and glycolytic ATP production rates, we found that MuSCs displayed a profound, >16-fold, increase in ATP production via the mitochondria during activation (Fig. 3E). As with OCR, MuSCs from juvenile mice displayed both the highest rates of mitochondrial ATP production and the most rapid increase (Fig. 3E). MuSCs from adult animals showed significantly higher mitochondrial ATP production rates than MuSCs from aged mice. Interestingly, we did not find any statistically significant increases in glycolytic ATP production rates during activation in MuSCs from any age group (Fig. 3F). Our results show that upon activation, there is a tremendous increase in mitochondrial respiration. Moreover, as the rate and magnitude of this increase maps closely with the speed with which MuSCs progress into and through the cell cycle, our results suggest that mitochondrial respiration may be an important driver of early MuSC activation.

**Figure 3:**
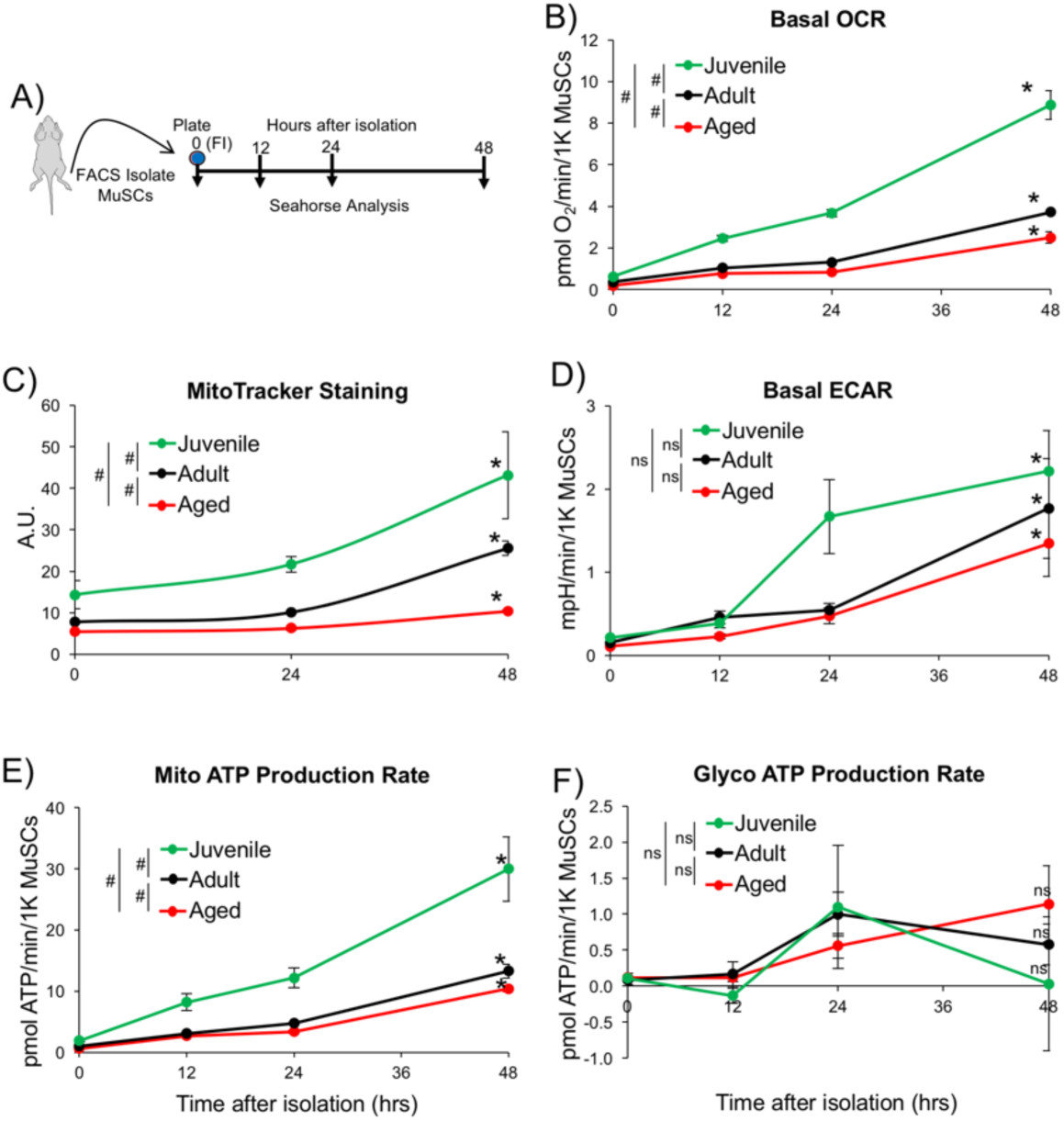
Age-associated defects in the induction of mitochondrial activity during MuSC activation. A) Schematic representation of experiments analyzing metabolic flux of MuSC following isolation. B) Quantification of cellular bioenergetics using seahorse extracellular flux assay. MuSCs isolated from juvenile (green line), adult (black line) and aged (red line) mice were cultured for 12 hours, 24 hours and 48 hours post-isolation *ex vivo* and subjected to OCR measurements using a Seahorse Bioanalyzer. Basal OCR of MuSCs through the course of activation presented as pmol O_2_/min/1K MuSCs. C) Quantification of MuSCs MTDR staining at 0 (FI), 24, and 48 hours after isolation. D) Basal ECAR presented as mpH/min/1K MuSCs. E) Mito ATP production rates and F) Glyco ATP production rate of MuSCs through the course of activation presented as pmol ATP/min/1K MuSCs. B-F) Each data point presented as mean ± SEM (N = 3-7). # denotes P < 0.01 comparing age groups by 2-way ANOVA. * denotes P < 0.01 comparing 48 hours vs 0 hour timepoints within an age group by Students t-test. ns denotes not statistically significant.

### G_Alert_ rescues age-associated changes in MuSC mitochondrial and activation function

We previously showed that the activation function of MuSCs could be enhanced by inducing them into a primed phase of quiescence, termed G_Alert_ ^11^. We and others have also shown that MuSCs, which were in G_Alert_ at the time which they were activated, required significantly less time to enter and complete the cell cycle and had greater muscle regenerative function than non-primed MuSCs in the G_0_ phase of quiescence^11–13^. We further showed that the transition of MuSCs into G_Alert_ correlated with modest but statistically significant increases in MTDR staining and cellular ATP levels as well as upregulation of transcriptional programs involved in mitochondrial oxidative phosphorylation in freshly isolated MuSCs ^11^. Consistent with previous work, using MuSCs isolated from muscles that were contralateral from the muscles in which we had induced an injury, as a model of G_Alert_^11^ (Fig. 4A), we found that freshly isolated G_Alert_ MuSCs had higher levels of MTDR staining than MuSCs isolated from animals that had not been subject to injury (non-injured, G_0_) (Fig. 4B). When cultured, a greater number of G_Alert_ MuSCs entered the cell cycle and incorporated EdU at 24 hours after isolation (Fig. 4C). Advancing on this, we also found that freshly isolated G_Alert_ MuSCs displayed greater OCR (Fig. 4D) and mitochondrial ATP production rates than MuSCs from non-injured animals of the same age (Fig. 4E). Interestingly, this trend of higher levels of OCR and mitochondrial ATP production rates continued through the course of activation (Fig. 4F-G). Combined, these data demonstrate that, when activated, G_Alert_ MuSCs possess higher mitochondrial function that correlates with their rapid entry into the cell cycle.

**Figure 4:**
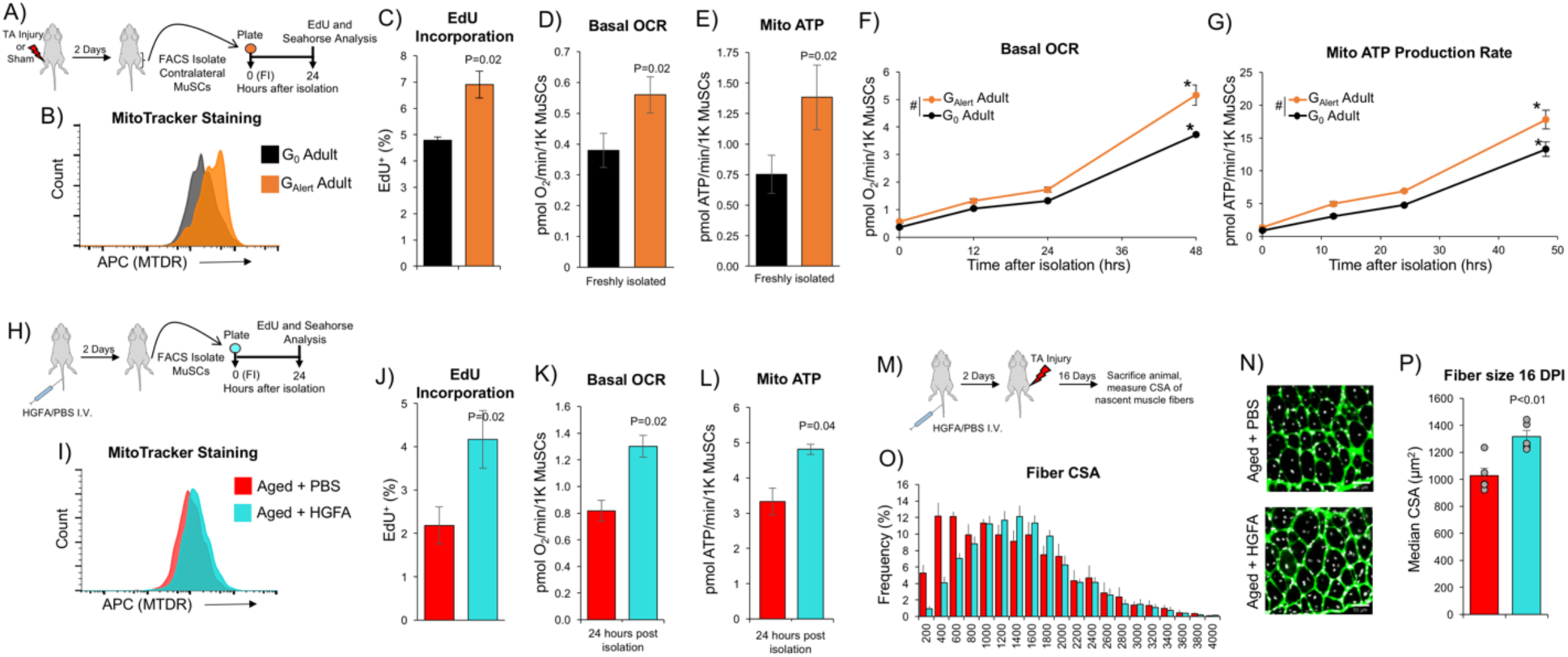
G_Alert_ rejuvenates MuSCs by increasing mitochondrial activity. A) Schematic representation of the experimental procedure and location of the G_Alert_ MuSCs relative, contralateral, to the injury site. B) Representative FACS MTDR histogram of freshly isolated MuSCs from adult mice contralateral to the site of an injury (G_Alert_-orange) and sham injured, non-injured, mice (G_0_-black). C) Quantification of EdU incorporation in G_Alert_ and G_0_ adult MuSCs cultured *ex vivo* for 24 hours. Data presented as mean ± SEM (N = 3-4) and significance was calculated by student’s unpaired t-test. D) Quantification of OCR in freshly isolated G_Alert_ and G_0_ adult MuSCs presented as pmol O_2_/min/1K MuSCs. E) Mito ATP production rates in freshly isolated G_Alert_ and G_0_ adult MuSCs presented as pmol ATP/min/1K MuSCs. F) Basal OCR and G) Mito ATP production rate for G_Alert_ and G_0_ adult MuSCs at 12, 24, and 48 hours after isolation and presented in a time-course curve. Values obtained from the quantification of freshly isolated (time 0) MuSCs (Fig. 4D-E) were used to plot the time-course curves. Each data point presented as mean ± SEM (N = 3-7). # denotes P < 0.01 comparing G_Alert_ vs. G_0_ by 2-way ANOVA. * denotes P < 0.01 comparing 48 hours vs 0 hour timepoints within each group by Students t-test. H) Schematic representation of the experimental procedure of HGFA-induced G_Alert_ MuSCs in aged mice. HGFA or PBS vehicle was administered through intravenous (I.V.) tail vein injections. I) Representative FACS MTDR histogram of freshly isolated MuSCs from HGFA injected aged mice (turquoise) and PBS injected aged mice (red). J) Quantification of EdU incorporation experiments of MuSC from aged animals injected with HGFA PBS and cultured for 24 hours in the presence of EdU. Data presented as mean ± SEM (N = 4-5) and significance was calculated by Student’s unpaired t-test. K) Basal OCR and L) Mito ATP production rate of MuSCs isolated from HGFA/PBS injected aged mice at 24 hours post-isolation. Data are presented as mean ± SEM (N = 3-7) and significance was calculated by Student’s unpaired t-test. M) Schematic representation of the experimental procedure for muscle regeneration assay. N) Representative IF-IHC staining of tibialis anterior (TA) sections from injured aged mice injected with HGFA or PBS (control). Laminin (Green) and DAPI (white). Scale bar 50μm. O) Histograms displaying the distribution of fiber CSA of the data that are summarized in Figure 4P. Each bar represents the mean ± SEM of the percentage of centrally nucleated fibers that fall within the bin range (N = 5). P) Median TA fiber CSA of injured control vs HGFA treated old mice (N = 5). Significance was calculated by Student’s t-test.

Next, we sought to determine if MuSCs from aged mice would display G_Alert_ mediated functional improvements similar to those of adult mice. To do this, we used systemic administration of Hepatocyte Growth Factor Activator (HGFA) as a pharmacologic model to induce MuSCs into G_Alert_ ^12^ (Fig. 4H). As expected, MuSCs isolated from aged animals two days after they had been administered HGFA had higher levels of MTDR staining (Fig. 4I) and entered the cell cycle more rapidly than MuSCs from animals that were administered PBS (control), as measured by EdU incorporation (Fig. 4J). MuSCs isolated from aged animals that had been administered HGFA also displayed higher levels of OCR and mitochondrial ATP production rates (Fig. 4K-L), similar to adult G_Alert_ MuSCs. Intriguingly, we found that aged animals that had been administered HGFA displayed significantly larger muscle fibers 16 days after muscle injury than control animals (Fig. 4M-P), consistent with our previous work in adult animals^12^. These results show that HGFA increases the mitochondrial function and activation speed of MuSCs in aged mice and correlates with improvements in muscle regeneration.

Taken as a whole, the data presented here demonstrate a tight correlation between MuSC mitochondrial, activation, and muscle regenerative functions - with age, there is a decrease in these MuSC functions, while conditions that induce MuSC into G_Alert_ induce a rejuvenation of these MuSC functions. Moreover, as we can identify changes in MuSC mitochondrial function immediately upon FACS isolation, many days before cell cycle entry and completion and muscle repair, this suggests that regulation of MuSC mitochondrial function may be an underlying driver of MuSC activation and muscle regenerative functions.

### Mitochondrial metabolic flux increases during MuSC activation

To gain deeper insights into the metabolic properties of MuSCs, we performed metabolomic analyses. Utilizing juvenile MuSCs, collected at four different time points - freshly sorted (0 hour), 12 hours, 24 hours and 48 hours post isolation (Fig. 5A). We performed targeted LC-MS metabolite analysis and were able to detect 143 metabolites at greater than seven-fold signal to noise ratio (Supplemental Data File). Principal component analysis (PCA) of the metabolite profiles revealed three distinct clusters ordered along PC axis 1; freshly isolated MuSCs, 12 hours post isolation, and a cluster that contains 24 and 48 hours after isolation (Fig. 5B). Similarly, spearman rank hierarchical clustering of the metabolite profiles, also shows three distinct clusters of metabolites that change during MuSC activation (Fig. 5C). The first cluster contains metabolites that are highly abundant in freshly isolated MuSCs and decline during activation, this cluster is enriched for metabolites in aminoacyl-tRNA biosynthesis, valine, leucine and isoleucine, and glycine, serine and threonine metabolic processes (Fig. S3). Cluster 2 is composed of metabolites that reach maximum abundance 12 hours after isolation and then decline at 24 and 48 hours. Cluster 2 is enriched for metabolites in propanoate and phenylalanine metabolism (Fig. S3). Cluster 3 contains metabolites that have low abundance in freshly isolated and 12 hour time points but are high in abundance at 24 and 48 hour time points. Cluster 3 is enriched for metabolites in nucleotide, proline and glutathione metabolism (Fig. S3). Overall, the metabolite profiles reveal a pattern of cells transitioning from a low energy state characterized by a relatively high AMP/ATP ratio to a high energy state with higher levels of oxidative metabolism as indicated by a dramatic increase in NAD/NADH ratios (Fig. 5D-E). Of note, of the metabolites that displays the most dramatic changes in MuSCs during activation is pyruvate. Intracellular pyruvate levels are high in freshly isolated MuSCs, and then display a tremendous ∼40-fold decrease in abundance during activation (Fig. 5F).

**Figure 5:**
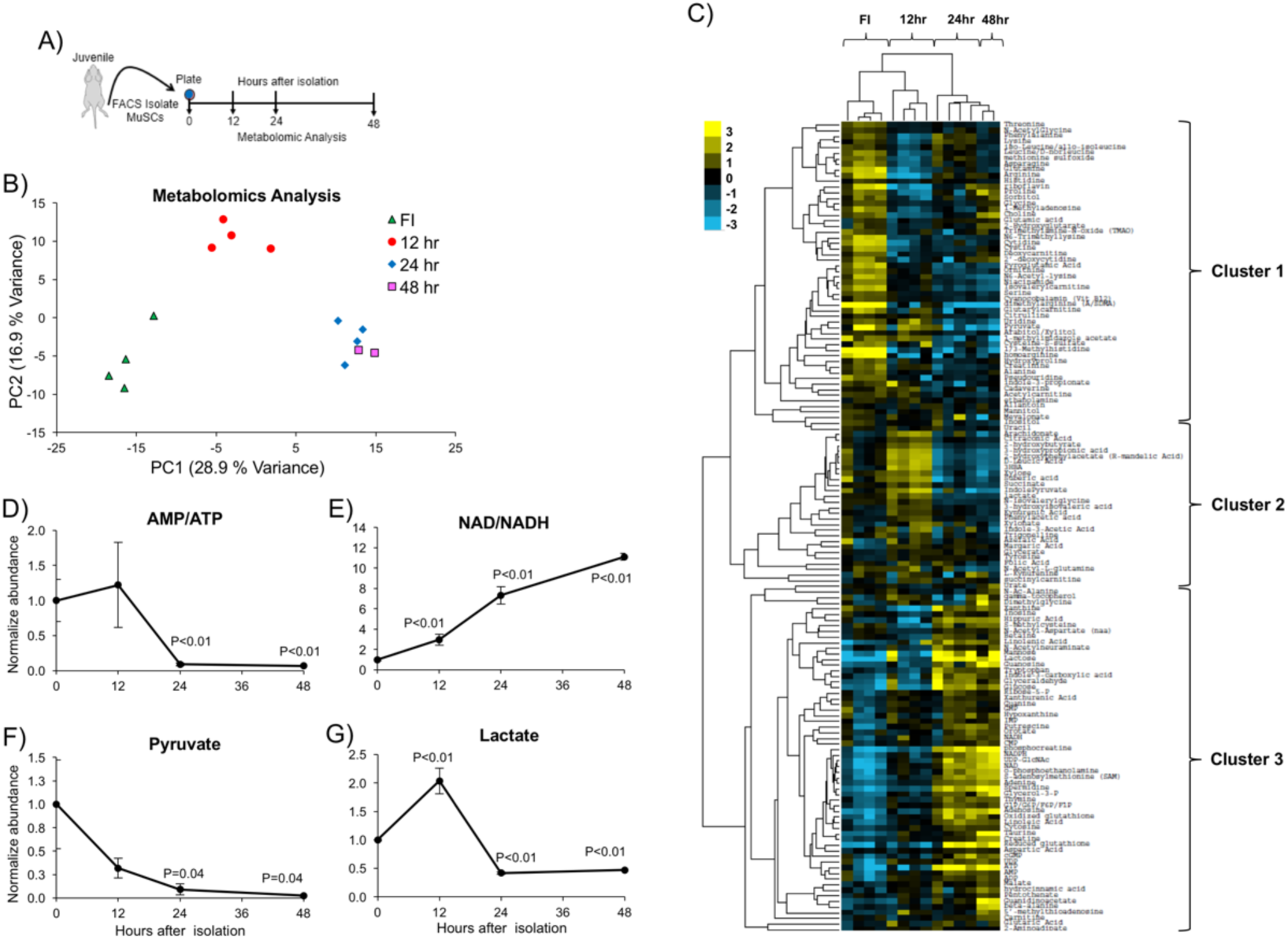
Mitochondrial flux increases during activation in MuSCs. A) Schematic representation of the experimental procedure to collect MuSCs for targeted metabolomics assay. B) PCA analysis of the metabolite profiles of MuSCs isolated from juvenile mice at different time-points post isolation- freshly isolated (FI) (N = 4), 12 hours (N = 4), 24 hours (N = 4) and 48 hours (N = 2). C) Heat map depicting normalized abundance of metabolites and spearman rank hierarchical clustering of the metabolite profiles revealed 3 distinct enrichment clusters of metabolites which vary as a function of time after isolation of MuSCs. D) Bar graph depicting the normalized AMP/ATP ratios calculated from relative metabolite abundance. E) Bar graph depicting the normalized NAD/NADH ratios calculated from relative metabolite abundance. F) Bar graph depicting the normalized pyruvate abundance. G) Bar graph depicting the normalized lactate abundance. D-G) Data are presented as mean ± SEM (N = 2-4) and P values represent the significance compared to FI (0 time point) and were calculated by ANOVA.

Pyruvate is a product of glycolysis and the two main pathways by which pyruvate is consumed in cells are: 1) reduction to lactate via LDHA and 2) oxidation to acetyl-CoA in the mitochondria via PDH. To test the role of LDHA in MuSC activation we utilized a conditional allele of LDHA combined with a MuSC-specific Pax7^CreER^ driver to ablate *Ldha* in MuSCs (LDHA-KO). We found that freshly isolated MuSCs from LDHA-KO mice displayed higher levels of MTDR staining than MuSCs from control mice (Fig. S4A-B). We also found that MuSCs from LDHA-KO animals had higher rates of mitochondrial ATP production (Fig. S4C). Moreover, after 40 hours in culture, we observed that a higher fraction of MuSCs from LDHA-KO mice had incorporated EdU than control MuSCs (Fig. S4D). We observed a similar effect when we cultured MuSCs from aged mice in a selective inhibitor of LDHA, GSK283780A^46^ (herein termed GSK)(Fig. S4E). MuSCs from aged mice cultured in GSK showed increases in mitochondrial ATP production rates (Fig. S4F) and an increased ability to incorporate EdU in 48 hours (Fig. S4G). Combined, these results show that genetic and pharmacologic inhibition of LDHA improves aspects of MuSC activation and surprisingly, suggests that pyruvate metabolism via LDHA is not a critical metabolic pathway that is required for activation. Consistent with this, we found only modest changes in intracellular lactate during MuSC activation (Fig. 5G). Taken together, these data highlight the dramatic decrease in intracellular pyruvate (Fig. 5F), mirroring the massive increase in mitochondrial ATP production rate (Fig. 3E) and decrease in AMP/ATP ratio (Fig. 5D), with only modest changes in intracellular lactate (Fig. 5G) and ECAR (Fig. 3F). This along with the improvements in activation induced by inhibiting LDHA (Fig. S4D, S4G) suggest that pyruvate is oxidized by the mitochondria during activation.

### PDH activity correlates with MuSC activation and muscle regenerative function

PDH is the primary regulator of pyruvate oxidation in the mitochondria. PDH enzymatic activity is strongly inhibited by pyruvate dehydrogenase kinases (PDKs)-mediated phosphorylation. The results outlined above led us to hypothesize that PDH would be phosphorylated early in MuSC activation and dephosphorylated later in activation. Indeed, by analyzing phospho-Ser293-PDH (pPDH) levels by immunofluorescence we found a modest albeit statistically significant decrease in pPDH levels at 24 hours after isolation compared to freshly isolated MuSCs from juvenile mice (Fig. 6A). Moreover, we found that pPDH levels in freshly isolated MuSCs showed a direct inverse correlation with mitochondrial activity and speed of activation across all of the model systems we have tested. We found that pPDH levels show an age-associated increase in MuSCs (Fig. 6B); correlating with age-associated decreases in mitochondrial activity (Fig. 2B-E, 2G) and activation speed (Fig. 1J). Likewise, pPDH levels were lower in adult G_Alert_ MuSCs (Fig. 6C) and in MuSCs from aged mice treated with HGFA (Fig. 6D), both of which are models of elevated mitochondrial function and more rapid cell cycle entry and activation (Fig. 4). Combined, these data suggest that PDH activity regulates the speed of MuSC activation by limiting pyruvate entry into the mitochondria.

**Figure 6:**
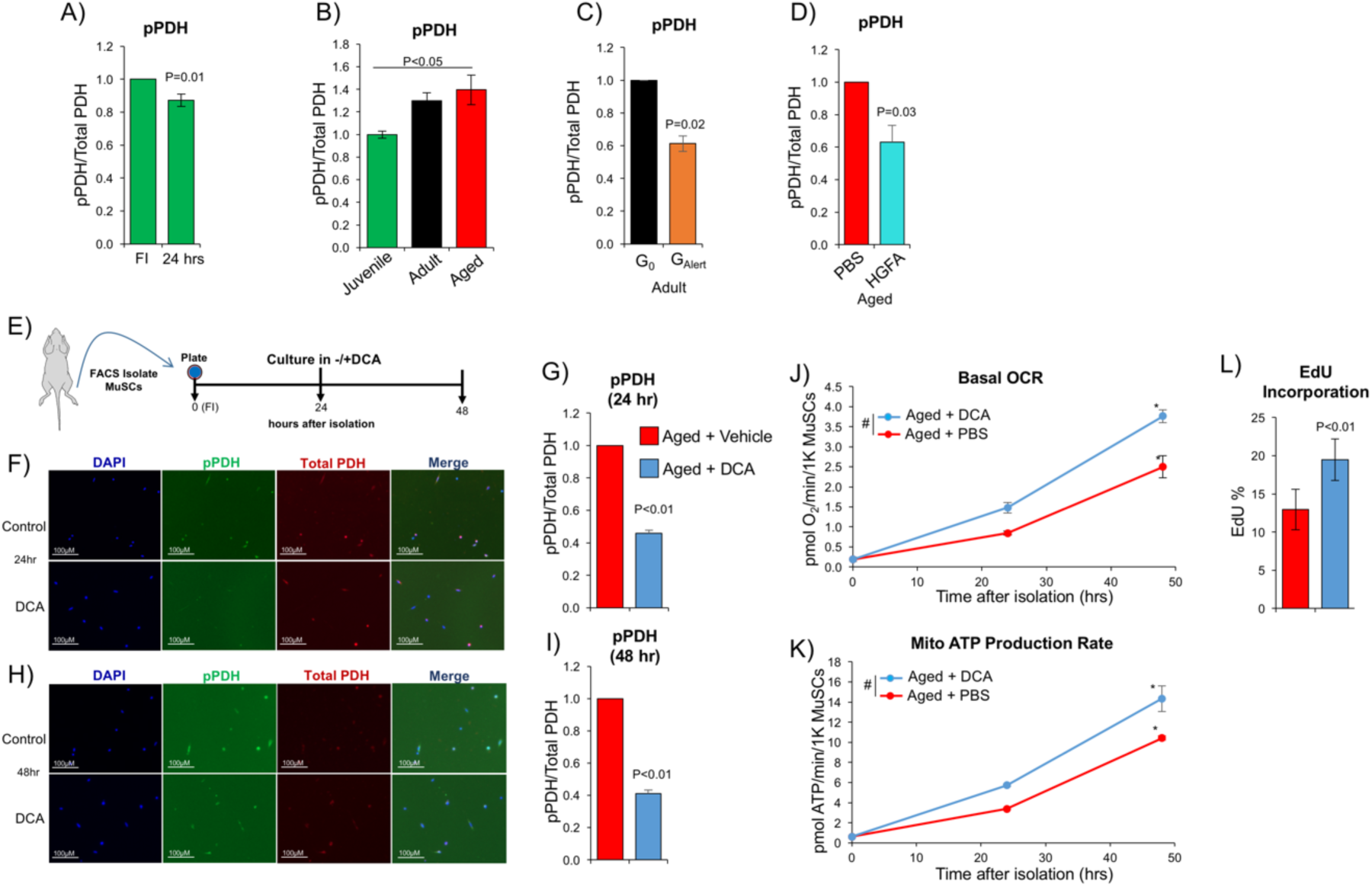
Depression of PDH promotes mito ATP production and MuSC activation. A) Quantification of pPDH of freshly isolated (FI) and 24 hours *ex vivo* cultured MuSCs isolated from juvenile mice. B) Quantification of pPDH of FI MuSCs from juvenile, adult and aged mice. C) Quantification of pPDH of FI MuSCs from non-injured adult mice (G0) and muscles contralateral to an injury in adult mice (G_Alert_). D) Quantification of pPDH of FI MuSCs from aged animals two days after injection with HGFA or PBS vehicle. E) Schematic representation of experimental procedure for dichloroacetate (DCA) treatment. F) Representative IF-ICC images of pPDH and PDH staining of control versus DCA treated MuSCs isolated from aged mice at 24 hours after isolation. G) Quantification of pPDH of MuSCs from aged mice cultured in DCA or PBS vehicle for 24 hours after isolation. H) Representative IF-ICC images of pPDH and PDH staining of control versus DCA treated MuSCs isolated from aged mice at 24 hours after isolation. I) Quantification of pPDH of MuSCs from aged mice cultured in DCA or PBS vehicle for 48 hours after isolation. J) Quantification of basal OCR and K) Mito ATP production rates of aged MuSCs cultured in DCA or vehicle for 24 and 48 hours after isolation. Each data point presented as mean ± SEM (n = 3-7). # denotes P < 0.01 comparing age groups by 2-way ANOVA. * denotes P < 0.01 comparing 48 hours vs 0 hour timepoints within an age group by Students t-test. L) Quantification of EdU incorporation in control vs DCA treated MuSCs isolated from aged mice at 48 hours post-isolation. Data are presented as mean ± SEM (n=3) and significance was calculated by student’s paired t-test. A-D, G, and I) Data are presented as the mean ± SEM. of the normalized ratio of pPDH signal to total PDH signal measured by IF analysis of MuSCs. Student’s t-test was used to calculate significance (N = 3-7).

To directly test this hypothesis, we utilized dichloroacetate (DCA), an inhibitor of PDKs, to activate PDH. Treating MuSCs from aged mice with DCA significantly reduced pPDH levels compared to control treated aged MuSCs (Fig. 6E-I), indicating that DCA is inhibiting PDKs. Importantly, this decrease in pPDH directly correlated with increases in mitochondrial activity (Fig. 6J-K). DCA treatment significantly increased the OCR and mitochondrial ATP production rates of MuSCs from aged mice (Fig. 6J-K). Finally, we found that a significantly greater number of MuSCs from aged mice cultured in DCA versus control conditions incorporated EdU in 48 hours (Fig. 6L). Take as a whole, these results show that pPDH levels strongly predict MuSC activation function and that reducing pPDH levels increases the speed of cell cycle entry for aged MuSCs.

Next, we sought to determine if these DCA-mediated improvements in MuSC function could be translated *in vivo*. To do this, we performed crush-induced injuries of TA muscle in aged mice and then administered either DCA or PBS to animals subcutaneously over the site of muscle injury (Fig. 7A). Thirty hours after injury, we found that a greater number of MyoD^+^ cells had incorporated EdU in the injured muscles of animals that had been administered DCA than control animals (Fig. 7B), suggesting that DCA treatment increased the speed of cell cycle entry. Taking these experiments a step further, we measured the effect of DCA on muscle regeneration in aged mice. Using a similar experimental setup, we continued DCA or PBS control administration for the first 3 days after a crush injury and then euthanized animals on day 8 after injury and assessed the size of centrally nucleated fibers by morphometric analysis (Fig. 7C). Strikingly, we found that regenerating fibers in animals that had been administered DCA had a significantly larger CSA than animals that received PBS control (Fig. 7D-F). These data show that DCA improves aspects of MuSC activation and muscle regenerative functions in the context of age-associated impairments. Overall, these data strongly suggest a model in which age-associated changes in MuSC metabolic function, via PDH activity, directly contribute to decline in activation and regeneration functions. Moreover, they also suggest that interventions to increase MuSC mitochondrial function, via inducing MuSCs into G_Alert_ and/or by suppressing PDK, may be a therapeutic approach to rejuvenate the function of MuSCs in aging.

**Figure 7:**
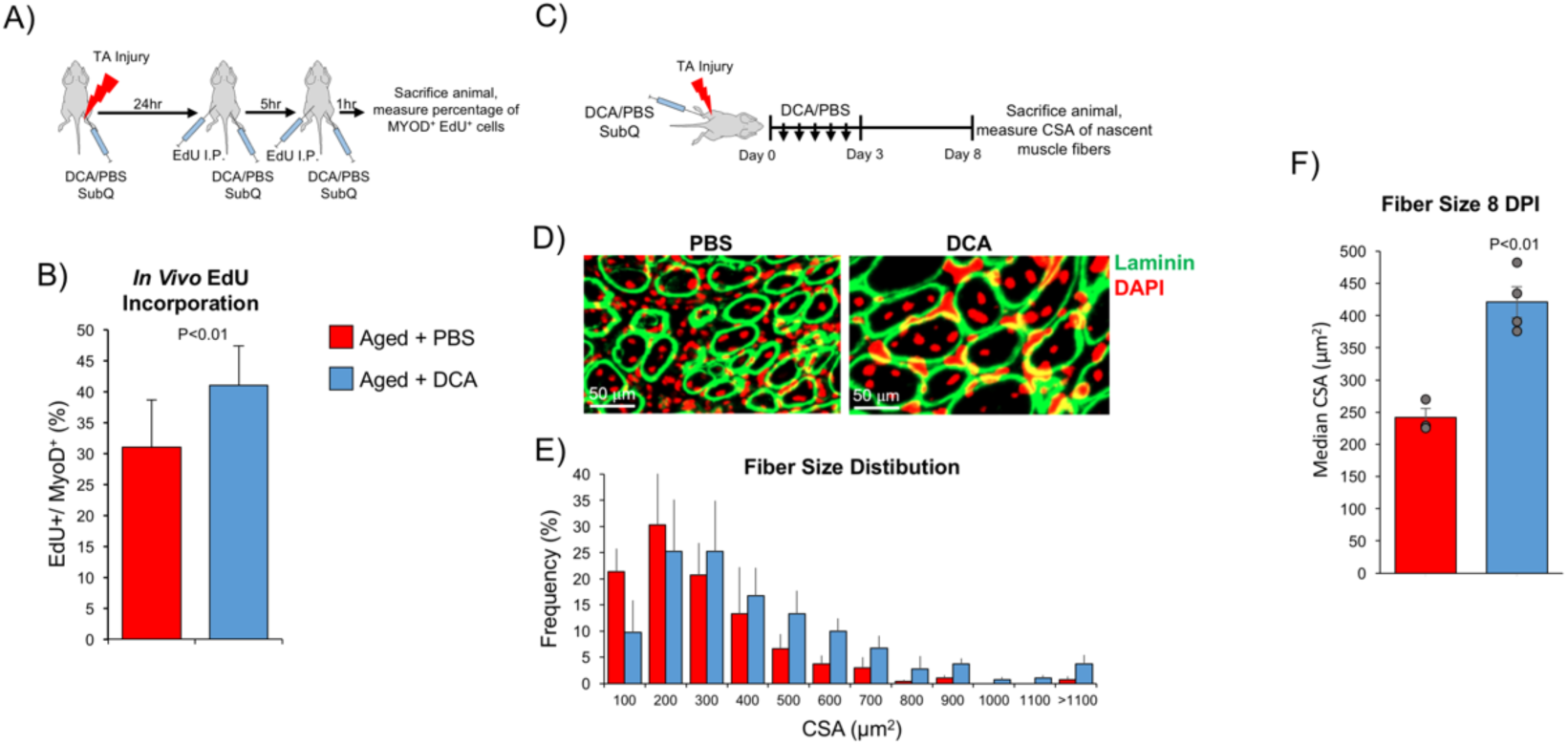
DCA promotes MuSC activation and regenerative functions *in vivo*. A) Schematic representation of the experimental procedure to analyze *in vivo* EdU incorporation follow muscle injury and DCA administration. B) Quantification of the percentage of Edu^+^ MyoD^+^ cells in the injured TAs of aged animals admistered DCA or vehicle (PBS). EdU incorporation and MyoD were analyzed by IF-IHC staining of muscle cryosections. Data are presented as mean ± SEM (N = 3) and significance was calculated by Student’s paired t-test. C) Schematic representation of the experimental procedure to analyze the effect of DCA on muscle regeneration in aged mice. D) Representative image of IF-IHC staining of tibialis anterior (TA) sections from control and DCA treated injured aged mice at 8 days post-injury (DPI). Laminin (green) and DAPI (red). E) Histogram displaying the size distribution of centrally nucleated fibers 8 DPI from aged animals administered DCA or vehicle 8 DPI. Each bar represents the mean ± SEM of the percentage of centrally nucleated fibers that fall within the bin range (N = 4). Fiber CSA was determined by morphometric analysis of centrally nucleated fibers in TA cryosections. D) Median fiber CSA of injured aged mice treated with DCA or PBS 8DPI. Data a presented as mean ± SEM (N = 4) and significance was calculated by Student’s t-test.

## DISCUSSION

Here, we provide a detailed analysis of MuSC metabolism and mitochondrial function during the transition from a quiescent state into the cell cycle and have implicated PDH-regulated mitochondrial flux as a critical pathway that contributes to age-associated defects in MuSC activation. We find that with age, there is a marked decrease in the speed by which MuSC transit from G0-G1 to S phase and that this transition is the rate-limiting step that controls the speed of MuSC activation. We present data, using numerous physiologic, genetic, and pharmacologic *in vivo* models, to show that the mitochondrial function of freshly isolated MuSCs directly correlates with their activation and muscle regenerative functions. This suggests that the mitochondrial activity and respiration levels in quiescent or freshly isolated MuSCs could be used as an early prognostic factor for muscle regenerative function.

Our findings support a mechanistic model whereby, PDH controls the speed of MuSC activation by controlling flux of pyruvate into the mitochondrial. We found that activated MuSCs display a dramatic, >16-fold, increase in mitochondrial activity and function, which correlates with a 40-fold drop in intracellular pyruvate levels and decreased PDH phosphorylation. Further, we find that pharmacological means to reduce PDH phosphorylation (increasing PDH activity) are sufficient to drive more rapid induction in mitochondrial function during activation, resulting in a more rapid entry into the cell cycle. Importantly, we show that we can utilize this pathway *in vivo* to improve MuSC activation and muscle regenerative functions.

PDH-regulated flux of pyruvate into the TCA cycle has been shown to be critical for MuSC differentiation and tissue regeneration ^28^. Defective MuSC mitochondrial function results in impaired tissue regeneration ^29^ and MuSCs with greater mitochondrial content and oxidative phosphorylation are associated with increased capability of initiating myogenic colonies ^22^. Interestingly, across tissue-specific stem cells there is increasing evidence for a link between stem cell metabolism and epigenetics ^25,26,33,47^. Quiescent HSCs rely on glycolysis during self-renewal state. However, upon activation their metabolism shifts from glycolysis to mitochondrial respiration ^33^. Neural stem cells also exhibit a similar metabolic switch during their transition from quiescence to differentiation ^31^. Hair follicle stem cells maintain a predominantly glycolytic metabolic state to control their self-renewal capacity ^30^. Promoting anaerobic glycolysis accelerates hair follicle stem cell activation and the hair cycle. Overall, our current data are consistent with emerging work in MuSCs and many other tissue-specific stem cell models showing that metabolism plays an important role in controlling stem cell fate and function.

We have previously described G_Alert_ as an adaptive response, whereby in response to systemic factors, MuSCs are induced to transition into a primed/poised state of quiescence, in an mTORC1-dependent manner. MuSC activation and muscle regenerative functions could be improved so long as MuSCs were in G_Alert_ at the time of activation (i.e. injury). Our current work suggests an underlying mechanism by which G_Alert_ MuSCs possess improved functional potential; by derepression of PDH and increased mitochondrial flux. Here, we examined two different models of alert cells, physiologic (injury-induced) and pharmacologic (HGFA-induced). Both resulted in decreased levels of PDH phosphorylation. We advance this work, showing that inhibiting PDK (and activating PDH) in *ex vivo* or *in vivo* systems after MuSC activation are sufficient to recapitulate the functional and biologic effects of G_Alert_. Further work needs to be done to test necessity of the PDK-PDH pathway in the G_Alert_ response.

DCA has been the subject of several experimental and clinical studies for the treatment of lactic acidosis and for investigating its impact on substrate level phosphorylation in skeletal muscle during transition from resting state to moderate-intensity exercise ^48,49^. These studies suggest that DCA-mediated PDH activation leads to decrease in lactate production in skeletal muscle. DCA has also been tested to show that increased availability of acetyl-CoA at the start of exercise decreases reliance on glycogenolysis and substrate level phosphorylation from phosphocreatine ^50,51^. While there is much evidence showing the effect of DCA on muscle metabolism and function, this is the first report, to our knowledge, of DCA having a direct impact on muscle repair/regeneration. Our data suggest that DCA may have a therapeutic application on improving muscle repair and regeneration in the context of aging.

## EXPERIMENTAL METHODS

### Mice

All the experiments were performed with male C57BL/6 mice of three different age groups-juvenile 1.5-2 months-old, adult 4-6 months-old and aged 24-30 months-old (Charles River), except for experiments using conditional LDHA-KO mice. Animals harboring conditional LDHA alleles (Jax # 030112), were bred with bred with the tamoxifen sensitive Pax7^CreER^ driver ^52^ and the Rosa26-stop-flox-TdTomato alleles (Jax #007905). LDHA-KO mice were genotyped by PCR of tail DNA. Tamoxifen (Sigma) was prepared in corn oil and administered daily at 150 mg/kg body weight via intraperitoneal injection over 5 days. One week before the experiment, mice were transferred to the Animal facility at the University of Southern California (USC). All animal protocols were approved by the Institutional Animal Care and Use Committee (IACUC) board of USC.

### HGFA Injections

We used purified, recombinant, active HGFA (R&D Systems) (R&D Systems Cat# 1200-SE; Lot # NNJ011502A) to administer aged mice via intravenous tail vein injection at a dose of 1 µg diluted into 200µL of sterile PBS. Control mice were administered with 200µL of sterile PBS alone.

### Tibialis Anterior Injury

Tibialis Anterior (TA) muscle injuries were induced by crush injury to the TA or by intramuscular injection with 70µL of a 1.2% solution of BaCl_2_ as previously described ^12^. For crush injuries, animals were anesthetized using isoflurane, a small incision was made along the length of a TA muscle, and the TA was mechanically crushed using a sterilized hemostat. The skin was then sutured and animals were administered buprenorphine and Baytril.

### MuSC isolation and culture

MuSCs were isolated using fluorescence activated cell sorting (FACS) instrumentation (BD FACSAria IIIu) as described previously ^54^. Briefly, hindlimb muscles were collected and enzymatically digest with collagenase and dispase. MuSCs were isolated as a population of CD45^−^; CD31^−^; Sca-1^−^; α-integrin^+^; VCAM^+^ cells into plating medium (Ham’s F-10 (Cellgro) containing 5ng/mL basic fibroblast growth factor (Invitrogen), 10% FBS (Invitrogen) and 1x penicillin/streptomycin (GIBCO)). Freshly isolated MuSCs were then plated onto poly-D-lysine (Millipore) and ECM (Sigma E1270) coated 8-well chamber slides (Lab-Tech II). For cells cultured beyond 12 hours, the medium was replaced with culturing medium (Ham’s F-10 (Cellgro) containing 10% horse serum (Invitrogen), 10% FBS (Invitrogen), and 1x penicillin/streptomycin (GIBCO)).

### Dichloroacetate Injections

We administered DCA (Sigma Cat# 347795) to aged mice via subcutaneous route over the injured TA region at a dose of 5mg diluted into 100uL of sterile PBS. For muscle regeneration experiments, we administered two subcutaneous injections of DCA every day at an interval of 8 hours for the first three days. For *in vivo* EdU incorporation experiments, we administered a total of three subsequent subcutaneous DCA injections-at the time of injury, 24 hours after injury, and 29 hours after the injury. Similarly, control mice were injected with 100uL sterile PBS alone.

### EdU Incorporation Assay

For *ex vivo* experiments, isolated MuSCs were cultured in EdU containing medium at a final concentration of 10 μM and replenished every 14 hours until cells were fixed. For *in vivo* EdU incorporation assay, mice were injected with 100 μL of 10mM EdU diluted into 200ul of sterile PBS at 24 hours after injury and 29 hours after injury. EdU was detected in fixed cells or tissue sections using the Click-It Kit (Invitrogen) according to the manufacturer’s instructions. *In vitro* EdU incorporation data are presented as the percentage of total cells (quantified by DAPI). *In vivo* EdU data are presented as the percentage of MyoD positive cells that possessed detectable EdU signal.

### Time-lapse microscopy

Time-lapse microscopy was performed using Zeiss Axio Observer Z1 mounted with a 10X objective. Chamber slides with cultured cells were transferred to a temperature and CO2 controlled chamber mounted on the microscope stage. Time-lapse image acquisition was carried out every 10min and visualization was made using Axiovision software. The time to complete the first division after plating was analyzed for only cells present in the field of view during acquisition.

### Cell and tissue immunostaining

Immunocytochemistry was performed on cells cultured in 8-well chamber slides. Cells were fixed in 4% PFA for 10 min, followed by permeabilization in 0.3% Triton X-100 PBS solution for 20 min. Permeabilized cells were then blocked in donkey serum for 30 min. pPDH was detected using PhosphoDetect Anti-PDH-E1α (pSer^293^) Rabbit pAb (EMD Millipore Cat# AP102) primary antibody at a dilution of 1:100 and secondary detection was performed using donkey anti-rabbit Alexa 488 antibodies (Invitrogen) at a dilution of 1:500. Total PDH was detected using Anti-Pyruvate Dehydrogenase E1-alpha subunit antibody (Abcam Cat# ab110330) primary antibody at a dilution of 1:100 and secondary detection was performed using donkey anti-mouse Alexa 647 antibodies (Invitrogen). pRB staining was performed following EdU staining by first permeabilizing the cells and then blocking in donkey serum for 20 min. After blocking, the cells were stained using Phospho-Rb (Ser807/811) (D20B12) XP® Rabbit mAb (Cell Signaling Technology) primary antibodies at a dilution of 1:100. Secondary staining was performed using donkey anti-rabbit Alexa 488 antibodies (Invitrogen) at a dilution of 1:500. All the antibodies were diluted in a solution of PBS, 10% donkey serum and 0.3% Triton X-100. Immunofluorescence immunohistochemistry (IF-IHC) was performed on TA muscle sections dissected immediately after euthanasia. After dissection, the tissue was mounted with tragacanth gum and snap-frozen in liquid nitrogen cooled isopentane. Mounted tissue was cryo-sectioned into 8μm sections and fixed in 4% PFA for 10 min, followed by washing with 0.3% Triton X-100 PBS solution. For muscle fiber size measurement, sections were stained with laminin antibodies (Santa Cruz sc-59854) at a dilution of 1:500 and secondary detection was performed using goat anti-rat Alexa 488 antibodies (Invitrogen). Fiber area was measured by using ImageJ software. MyoD staining was performed with the Zenon Mouse IgG Labeling Kit (Invitrogen) according to the manufacturer’s instructions. MyoD antibodies (BD Cat#554130) were used at a 6:1 molar ratio of Fab to MyoD antibody. All the antibodies were diluted in a solution of PBS, 10% donkey serum and 0.3% Triton X-100.

### FLIM

Freshly isolated MuSCs from juvenile, adult and aged mice were plated in an 8-well slide and incubated for 30 min before the imaging. A Zeiss LSM-780 inverted microscope with a live cell work station (37C, 5% CO2) was employed for FLIM. Samples were excited using a 740 nm two photon laser and emission filters of 460/80 nm and 540/50 nm were used to acquire NAD(P)H and FAD signals, respectively. Each image was acquired with 256 * 256 pixel resolution and pixel dwelling was 12.41 μsec. Frame repetition was set to 20 and the readout was averaged. For imaging analysis, FAD intensity was used to generate a mask for mitochondria. Enzyme-bound NAD(P)H fraction (αbound, defined by the ratio of enzyme-bound NAD(P)H over total NAD(P)H) in mitochondria was then calculated using a previously reported phasor approach ^41^.

### Metabolomics

Freshly isolated MuSCs from juvenile mice were cultured for various time-points before trypsinizing them for LC-MS sample preparation. At respective time-points, MuSCs were trypsinized for 2 minutes at 37 degrees, followed by washing in PBS and centrifuging at 500g for 2 min at room temperature. After centrifugation, supernatant was discarded and the pellet was resuspended in PBS. 250,000 cells were collected for each sample and pelleted down by centrifugation at 500g for 5 min at room temperature. The supernatant was discarded and the pellet was resuspended in 1 ml 80:20 methanol:water solution and vortexed for 10 sec. Samples were then stored at −20 degrees for 30 minutes. Next, samples were sonicated in an ice bath for 10 min followed by vortexing for 10 sec. The samples were then centrifuged at 14000 g for 15 min at 4 degrees. Following centrifugation, 600ul of the supernatant was transferred to a new tube and dried using Speedvac at 30 degrees. The dried pellets were stored at −80 degree until shipping it to University of Washington Nathan Shock Metabolomics Core, in collaboration with the Northwest Metabolomics Research Center, for targeted metabolomic analysis. Statistical analysis was carried out using R (version 3.6.0). Metabolite abundance data was log2 transformed. To control the injection volume variation, we performed a single internal standard (IS) normalization, where the positive mode detected metabolites were normalized to a 2C_13_-tyrosine isotope and negative mode detected metabolites were normalized to a 3C_13_-lactate isotope that were spiked into each sample. In order to further remove the systematic variation between samples, we performed a Cyclic LOESS normalization, which had good performance using a MS benchmark data set ^53^. This normalization step is implemented using normalizeCyclicLoess function from the limma R package ^54^. Metabolite data that is Missing Not At Random (MNAR) corresponds to censored missing values caused by metabolite abundances that are below the limits of quantification (LOQ) ^55^. We selected 143 metabolites with < 60% missingness and performed imputation using the stochastic minimal value approach that is designed to impute MNAR missing data. This was implemented using the impute.MinProb function in the imputeLCMD R package. In order to remove this unwanted variation in the data, we used the Bioconductor SVA package ^56^ to identify and estimate two surrogate variables (SVs), which we include as covariates in the subsequent analyses. We fit a linear model to the normalized and imputed metabolomic data using the Bioconductor limma package ^54^, with coefficient for time points and adjusting the estimated SVs as covariates in our model. The limma package uses empirical Bayes moderated statistics, which improves power by ‘borrowing strength’ between metabolites in order to moderate the residual variance ^57^. Metabolites were considered significant with a false discovery rate (FDR) < 0.1. Pathway analysis was performed using MetaboAnalyst^58^ with the full list of metabolites detectable in the targeted metabolomics analysis as the background list (Supplemental Data).

### Seahorse assay

Freshly isolated MuSCs were plated in ECM coated 8-well seahorse culture miniplates and cultured for various times points. MuSCs were sorted into plating media. For cells cultured beyond 12 hours, the media was changed to culture media. One day before the assay, seahorse sensor cartridge was prepared according to manufacturer’s instructions and incubated overnight in a CO2-free incubator at 37°C. On the day of assay, culture medium was changed to seahorse assay medium (Buffered XF Base medium (DMEM) containing, 5mM HEPES, 1mM pyruvate, 2mM L-glutamine (Life Technologies), and 10 mM glucose). Propidium iodide was added to all the cultured wells and control wells in the seahorse miniplates followed by imaging. These images were analyzed (based on propidium iodide staining) to determine the number of live cells in each well and normalize the bioenergetics rates. After imaging, cells were equilibrated at 37°C in a CO_2_-free incubator for 45 min before running the assay. Oligomycin was added each at a final concentration of 1 μM, Carbonyl cyanide-4-(trifluoromethoxy)phenylhydrazone (FCCP) was added at a final concentration of 2 μM, and Rotenone and Antimycin were added at a final concentration of 0.5 μM each after injection to the appropriate port of the sensor cartridge. To begin the assay, sensor cartridge was run on the Seahorse XFp Extracellular Flux Analyzer followed by the cell culture miniplate, and the data were analyzed by Excel. ATP production rates were calculated using the methodology outlined in Quantifying Cellular ATP Production Rate Using Agilent Seahorse XF Technology white paper document.

## ACKNOWLEDGEMENTS

This work has been funded by grants from the NIH (R00AG041764), AFAR (Junior Faculty Grant), and The Baxter Family Foundation (to J.T.R.), the NIH (AR060868 and AR061002) (to A.S.B.), the NSF (DMS1561814) and NIH (AG049494 and AG013280)(to D.P) and the Human Frontier Science Program (LT000781/2016-L) (to A.B.H). The metabolomic analysis was funded by the University of Washington’s Nathan Shock Center (P30 AG013280, PIs: Rabinovitch, P.S. and Kaeberlein, M.) and Center for Translational Muscle Research (P30AR074990, to D.R.). The authors would like to acknowledge Xuefeng Sun for assistance in the animal studies.

## REFERENCES

1. Collins, C. A. et al. Stem Cell Function, Self-Renewal, and Behavioral Heterogeneity of Cells from the Adult Muscle Satellite Cell Niche. Cell 122, 289–301 (2005).

2. Sambasivan, R. et al. Pax7-expressing satellite cells are indispensable for adult skeletal muscle regeneration. Development 138, 4333–4333 (2011).

3. Lepper, C., Partridge, T. A. & Fan, C.-M. An absolute requirement for Pax7-positive satellite cells in acute injury-induced skeletal muscle regeneration. Development 138, 3639–3646 (2011).

4. Schultz, E., Gibson, M. C. & Champion, T. Satellite cells are mitotically quiescent in mature mouse muscle: An EM and radioautographic study. Journal of Experimental Zoology 206, 451–456 (1978).

5. Conboy, I. M. & Rando, T. A. The Regulation of Notch Signaling Controls Satellite Cell Activation and Cell Fate Determination in Postnatal Myogenesis. Developmental Cell 3, 397–409 (2002).

6. Cooper, R. N. et al. In vivo satellite cell activation via Myf5 and MyoD in regenerating mouse skeletal muscle. J. Cell Sci. 112, 2895 (1999).

7. Brack, A. S. & Rando, T. A. Intrinsic Changes and Extrinsic Influences of Myogenic Stem Cell Function During Aging. Stem Cell Rev 3, 226–237 (2007).

8. Rocheteau, P., Gayraud-Morel, B., Siegl-Cachedenier, I., Blasco, M. A. & Tajbakhsh, S. A Subpopulation of Adult Skeletal Muscle Stem Cells Retains All Template DNA Strands after Cell Division. Cell 148, 112–125 (2012).

9. Der Vartanian, A. et al. PAX3 Confers Functional Heterogeneity in Skeletal Muscle Stem Cell Responses to Environmental Stress. Cell Stem Cell 24, 958-973.e9 (2019).

10. Scaramozza, A. et al. Lineage Tracing Reveals a Subset of Reserve Muscle Stem Cells Capable of Clonal Expansion under Stress. Cell Stem Cell 24, 944-957.e5 (2019).

11. Rodgers, J. T. et al. mTORC1 controls the adaptive transition of quiescent stem cells from G0 to GAlert. Nature 510, 393 (2014).

12. Rodgers, J. T., Schroeder, M. D., Ma, C. & Rando, T. A. HGFA Is an Injury-Regulated Systemic Factor that Induces the Transition of Stem Cells into GAlert. Cell Reports 479–486 (2017) doi: https://doi.org/10.1016/j.celrep.2017.03.066.

13. Lee, G. et al. Fully reduced HMGB1 accelerates the regeneration of multiple tissues by transitioning stem cells to GAlert. Proc. Natl. Acad. Sci. U.S.A. 115, E4463–E4472 (2018).

14. Jones, D. L. & Rando, T. A. Emerging models and paradigms for stem cell ageing. Nature Cell Biology 13, 506–512 (2011).

15. Liu, L. & Rando, T. A. Manifestations and mechanisms of stem cell aging. J. Cell Biol. 193, 257–266 (2011).

16. Sousa-Victor, P. et al. Geriatric muscle stem cells switch reversible quiescence into senescence. Nature 506, 316–321 (2014).

17. Cosgrove, B. D. et al. Rejuvenation of the muscle stem cell population restores strength to injured aged muscles. Nature Medicine 20, 255–264 (2014).

18. Brack, A. S., Bildsoe, H. & Hughes, S. M. Evidence that satellite cell decrement contributes to preferential decline in nuclear number from large fibres during murine age-related muscle atrophy. Journal of Cell Science 118, 4813–4821 (2005).

19. Sahin, E. & DePinho, R. A. Linking functional decline of telomeres, mitochondria and stem cells during ageing. Nature 464, 520–528 (2010).

20. Brack, A. S. & Muñoz-Cánoves, P. The ins and outs of muscle stem cell aging. Skeletal Muscle 6, 1 (2016).

21. Fulle, S. et al. Age-dependent imbalance of the antioxidative system in human satellite cells. Experimental Gerontology 40, 189–197 (2005).

22. Cerletti, M., Jang, Y. C., Finley, L. W. S., Haigis, M. C. & Wagers, A. J. Short-term calorie restriction enhances skeletal muscle stem cell function. Cell Stem Cell 10, 515–519 (2012).

23. Pala, F. et al. Distinct metabolic states govern skeletal muscle stem cell fates during prenatal and postnatal myogenesis. J Cell Sci 131, (2018).

24. Zhang, H. et al. NAD+ repletion improves mitochondrial and stem cell function and enhances life span in mice. Science 352, 1436–1443 (2016).

25. Ryall, J. G. et al. The NAD+-Dependent SIRT1 Deacetylase Translates a Metabolic Switch into Regulatory Epigenetics in Skeletal Muscle Stem Cells. Cell Stem Cell 16, 171–183 (2015).

26. Yucel, N. et al. Glucose Metabolism Drives Histone Acetylation Landscape Transitions that Dictate Muscle Stem Cell Function. Cell Reports 27, 3939-3955.e6 (2019).

27. Csete, M. et al. Oxygen-mediated regulation of skeletal muscle satellite cell proliferation and adipogenesis in culture. Journal of Cellular Physiology 189, 189–196 (2001).

28. Hori, S., Hiramuki, Y., Nishimura, D., Sato, F. & Sehara-Fujisawa, A. PDH-mediated metabolic flow is critical for skeletal muscle stem cell differentiation and myotube formation during regeneration in mice. The FASEB Journal 33, 8094–8109 (2019).

29. Theret, M. et al. AMPKα1-LDH pathway regulates muscle stem cell self-renewal by controlling metabolic homeostasis. The EMBO Journal 36, 1946–1962 (2017).

30. Flores, A. et al. Lactate dehydrogenase activity drives hair follicle stem cell activation. Nature Cell Biology 19, 1017–1026 (2017).

31. Knobloch, M. et al. A Fatty Acid Oxidation-Dependent Metabolic Shift Regulates Adult Neural Stem Cell Activity. Cell Reports 20, 2144–2155 (2017).

32. Simsek, T. et al. The Distinct Metabolic Profile of Hematopoietic Stem Cells Reflects Their Location in a Hypoxic Niche. Cell Stem Cell 7, 380–390 (2010).

33. Takubo, K. et al. Regulation of Glycolysis by Pdk Functions as a Metabolic Checkpoint for Cell Cycle Quiescence in Hematopoietic Stem Cells. Cell Stem Cell 12, 49–61 (2013).

34. Conboy, I. M., Conboy, M. J., Smythe, G. M. & Rando, T. A. Notch-mediated restoration of regenerative potential to aged muscle. Science 302, 1575–1577 (2003).

35. Charville, G. W. et al. Ex Vivo Expansion and In Vivo Self-Renewal of Human Muscle Stem Cells. Stem Cell Reports 5, 621–632 (2015).

36. Siegel, A. L., Kuhlmann, P. K. & Cornelison, D. D. W. Muscle satellite cell proliferation and association: new insights from myofiber time-lapse imaging. Skelet Muscle 1, 7 (2011).

37. Jones, R. G. et al. AMP-activated protein kinase induces a p53-dependent metabolic checkpoint. Mol. Cell 18, 283–293 (2005).

38. Zhang, S. et al. Cellular energy stress induces AMPK-mediated regulation of glioblastoma cell proliferation by PIKE-A phosphorylation. Cell Death & Disease 10, 1–13 (2019).

39. de Almeida, M. J., Luchsinger, L. L., Corrigan, D. J., Williams, L. J. & Snoeck, H.-W. Dye-Independent Methods Reveal Elevated Mitochondrial Mass in Hematopoietic Stem Cells. Cell Stem Cell 21, 725-729.e4 (2017).

40. Stringari, C. et al. Phasor approach to fluorescence lifetime microscopy distinguishes different metabolic states of germ cells in a live tissue. PNAS 108, 13582–13587 (2011).

41. Stringari, C. et al. Metabolic trajectory of cellular differentiation in small intestine by Phasor Fluorescence Lifetime Microscopy of NADH. Scientific Reports 2, 568 (2012).

42. Mookerjee, S. A., Gerencser, A. A., Nicholls, D. G. & Brand, M. D. Quantifying intracellular rates of glycolytic and oxidative ATP production and consumption using extracellular flux measurements. J. Biol. Chem. 292, 7189–7207 (2017).

43. Romero, N., Rogers, G., Neilson, A. & Dranka, B. P. White paper: Quantifying Cellular ATP Production Rate Using Agilent Seahorse XF Technology.

44. van Velthoven, C. T. J., de Morree, A., Egner, I. M., Brett, J. O. & Rando, T. A. Transcriptional profiling of quiescent muscle stem cells in vivo. Cell Rep 21, 1994–2004 (2017).

45. Machado, L. et al. In Situ Fixation Redefines Quiescence and Early Activation of Skeletal Muscle Stem Cells. Cell Rep 21, 1982–1993 (2017).

46. Billiard, J. et al. Quinoline 3-sulfonamides inhibit lactate dehydrogenase A and reverse aerobic glycolysis in cancer cells. Cancer Metab 1, 19 (2013).

47. Fawal, M.-A. & Davy, A. Impact of Metabolic Pathways and Epigenetics on Neural Stem Cells. Epigenet Insights 11, (2018).

48. Howlett, R. A., Heigenhauser, G. J. F., Hultman, E., Hollidge-Horvat, M. G. & Spriet, L. L. Effects of dichloroacetate infusion on human skeletal muscle metabolism at the onset of exercise. American Journal of Physiology-Endocrinology and Metabolism 277, E18–E25 (1999).

49. Stacpoole, P. W., Harman, E. M., Curry, S. H., Baumgartner, T. G. & Misbin, R. I. Treatment of Lactic Acidosis with Dichloroacetate. New England Journal of Medicine 309, 390–396 (1983).

50. Timmons, J. A. et al. Increased acetyl group availability enhances contractile function of canine skeletal muscle during ischemia. J. Clin. Invest. 97, 879–883 (1996).

51. Timmons, J. A. et al. Substrate availability limits human skeletal muscle oxidative ATP regeneration at the onset of ischemic exercise. J. Clin. Invest. 101, 79–85 (1998).

52. Nishijo, K. et al. Biomarker system for studying muscle, stem cells, and cancer in vivo. FASEB J. 23, 2681–2690 (2009).

53. Li, B. et al. NOREVA: normalization and evaluation of MS-based metabolomics data. Nucleic Acids Res 45, W162–W170 (2017).

54. Ritchie, M. et al. LIMMA powers differential expression analyses for RNA-sequencing and microarray studies. Nucleic acids research 43, (2015).

55. Wei, R. et al. Missing Value Imputation Approach for Mass Spectrometry-based Metabolomics Data. Scientific Reports 8, 1–10 (2018).

56. Leek, J. T. & Storey, J. D. Capturing Heterogeneity in Gene Expression Studies by Surrogate Variable Analysis. PLOS Genetics 3, e161 (2007).

57. Smyth, G. K. Linear models and empirical bayes methods for assessing differential expression in microarray experiments. Stat Appl Genet Mol Biol 3, Article3 (2004).

58. Chong, J. et al. MetaboAnalyst 4.0: towards more transparent and integrative metabolomics analysis. Nucleic Acids Res. 46, W486–W494 (2018).

