## Supplementary material for "PDH Mediated Mitochondrial Respiration Controls the Speed of Muscle Stem Cell Activation in Muscle Repair and Aging": Fig. S

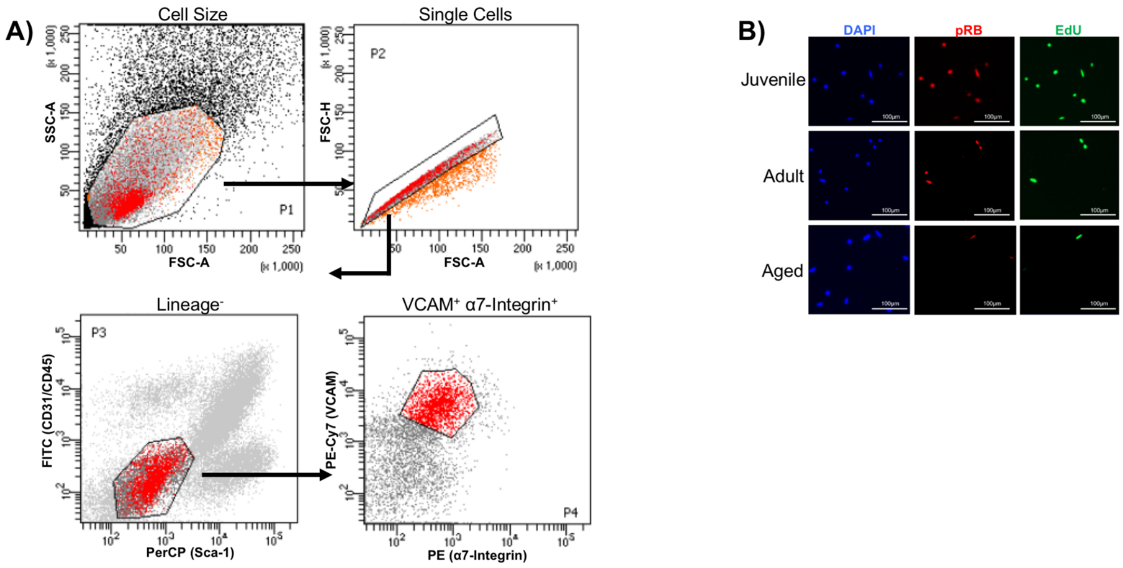

**Supplemental Figure 1: MuSC gating and pRB immunostaining**

A) Representative FACs plots and gating strategy to sort VCAM+ α-Integrin+ MuSCs from muscle tissue preparations. MuSCs stain negative for CD45/CD31/Sca1. B) Representative IF-ICC images of pRB and EdU stained MuSCs isolated from mice of different age groups fixed 40 hours post-isolation.

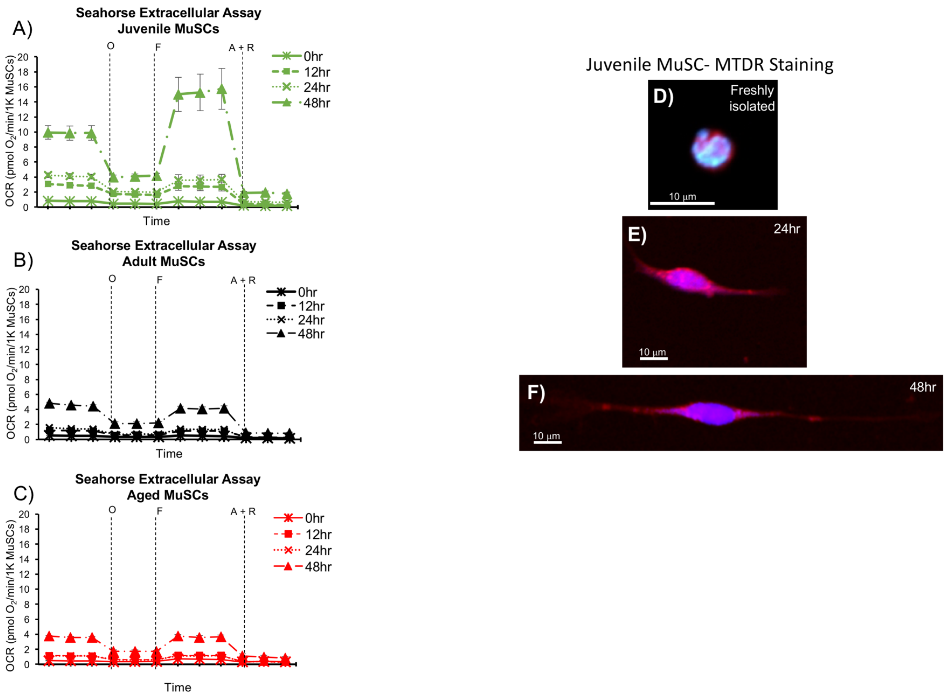

**Supplemental Figure 2: Seahorse OCR measurement plots and MTDR IF**

A-C) Representative OCR plots of MuSCs isolated from A) juvenile, B) adult, and C) aged mice cultured for various time-points post-isolation. Measurements were performed using a Seahorse XFp extracellular flux analyzer using mitochondrial stress kit. Each data point presented as mean $\pm$ SEM (N = 3-7).

D-F) Representative IF-ICC images of MitoTracker Deep Red (MTDR) stained MuSCs isolated from juvenile mice in freshly isolated, 24 hours, and 48 hours post-isolation time-points.

| | Pathway | Hits | Expected Hits | P Value | FDR |
|---|---|---|---|---|---|
| Cluster 1 | Aminoacyl-tRNA biosynthesis | 13 | 1.56 | 7.29E-10 | 6.12E-08 |
| | Valine, leucine and isoleucine biosynthesis | 3 | 0.26 | 1.62E-03 | 6.79E-02 |
| | Glycine, serine and threonine metabolism | 5 | 1.11 | 4.05E-03 | 1.13E-01 |
| | Pyrimidine metabolism | 5 | 1.27 | 7.41E-03 | 1.50E-01 |
| | Arginine biosynthesis | 3 | 0.46 | 9.16E-03 | 1.50E-01 |
| | Alanine, aspartate and glutamate metabolism | 4 | 0.91 | 1.14E-02 | 1.50E-01 |
| | Histidine metabolism | 3 | 0.52 | 1.35E-02 | 1.50E-01 |
| | D-Glutamine and D-glutamate metabolism | 2 | 0.20 | 1.43E-02 | 1.50E-01 |
| | Glyoxylate and dicarboxylate metabolism | 4 | 1.04 | 1.82E-02 | 1.70E-01 |
| | Arginine and proline metabolism | 4 | 1.24 | 3.24E-02 | 2.72E-01 |
| | Lysine degradation | 3 | 0.81 | 4.50E-02 | 3.44E-01 |
| Cluster 2 | Propanoate metabolism | 3 | 0.26 | 1.84E-03 | 1.55E-01 |
| | Phenylalanine metabolism | 2 | 0.14 | 7.41E-03 | 3.11E-01 |
| | Butanoate metabolism | 2 | 0.17 | 1.16E-02 | 3.24E-01 |
| | Phenylalanine, tyrosine and tryptophan biosynthesis | 1 | 0.05 | 4.44E-02 | 9.28E-01 |
| Cluster 3 | Purine metabolism | 15 | 2.59 | 7.58E-09 | 6.37E-07 |
| | Arginine and proline metabolism | 6 | 1.49 | 2.90E-03 | 1.07E-01 |
| | Glutathione metabolism | 5 | 1.10 | 3.82E-03 | 1.07E-01 |
| | Pyrimidine metabolism | 5 | 1.53 | 1.61E-02 | 3.19E-01 |
| | Galactose metabolism | 4 | 1.06 | 1.90E-02 | 3.19E-01 |
| | Pantothenate and CoA biosynthesis | 3 | 0.74 | 3.54E-02 | 4.83E-01 |
| | Glycine, serine and threonine metabolism | 4 | 1.33 | 4.10E-02 | 4.83E-01 |
| | beta-Alanine metabolism | 3 | 0.82 | 4.60E-02 | 4.83E-01 |

**Supplemental Figure 3: Pathway analysis on metabolites.**
Table summarizing the results of pathway enrichment analysis on the metabolites in the indicated cluster from Figure 5C. The ‘Pathway’ column indicates the metabolic pathway returned by MetaboAnalyst. The ‘Hits’ column indicates the number metabolites within the cluster that were in the annotated pathway. The ‘Expected Hits’ column indicates the number of hits expected based on the background list of metabolites included in the targeted metabolomic analyses. ‘P Value’ and False Discovery Rates (‘FDR’) are reported as calculated by MetaboAnalyst.

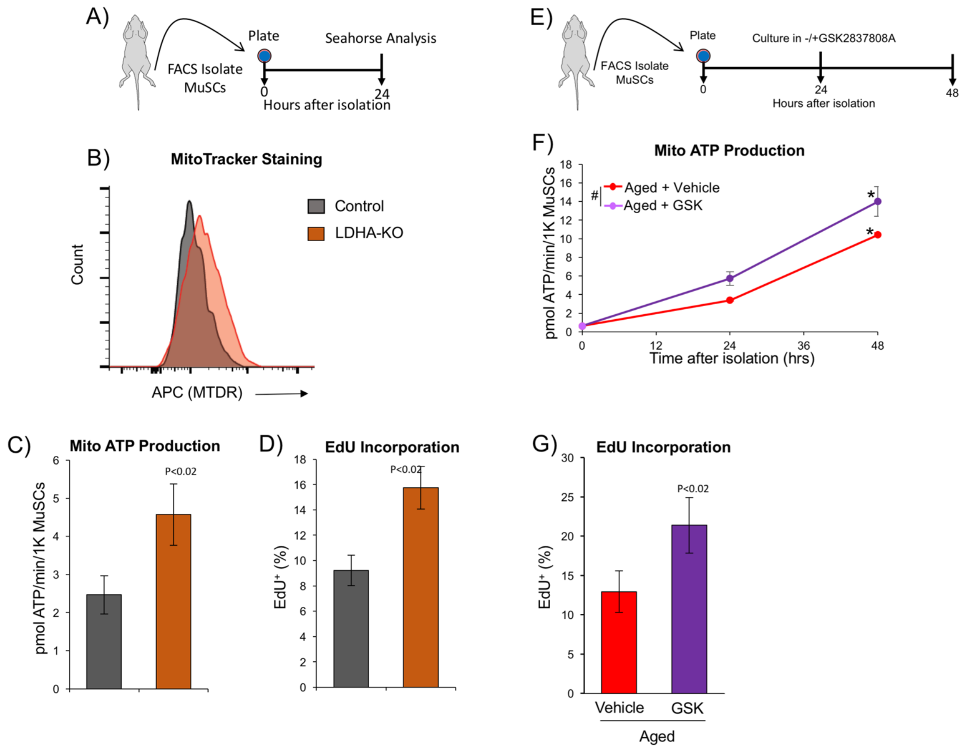

**Supplemental Figure 4: Inhibition of LDHA improves MuSC activation.**
A) Schematic representation of the experimental procedure for Seahorse Extracellular flux analysis of LDHA-KO MuSCs.
B) Representative FACS plot showing MTDR staining intensity of MuSCs from LDHA-KO and control.
C) Mito ATP production rate measurements for MuSCs isolated from LDHA control vs LDHA KO mice and cultured for 24 hours post-isolation Data presented as mean $\pm$ SEM (N = 4-5). Significance was calculated by Student's t-test.
D) Quantification of EdU incorporation in MuSCs isolated from LDHA control vs LDHA KO mice cultured *ex vivo* for 40 hours post-isolation. Data presented as mean $\pm$ SEM (n=4-5) and significance was calculated by Student's t-test.
E) Schematic representation of the experimental procedure test the effect of GSK treatment on MuSC activation.
F) Quantification of mito ATP production rate for MuSCs isolated from aged mice and treated with PBS vs GSK at 24 hours and 48 hours post-isolation in *ex vivo* culture, and presented in a time-course curve. Values obtained from the quantification of freshly isolated MuSCs were used to plot the time-course curves. # denotes P < 0.01 comparing GSK to vehicle treated by 2-way ANOVA. * denotes P < 0.01 comparing 48 hours vs 0 hour (FI) timepoints within an each group by Student's t-test.
G) Quantification of EdU incorporation in MuSCs isolated from aged mice and treated with vehicle (PBS) vs GSK during culture *ex vivo* for 48 hours post-isolation. Data presented as mean $\pm$ SEM (N = 3) and significance was calculated by Student's t-test.

**Supplemental Data File (.xlsx): Metabolomics data.**
Normalized_Counts_Log2 tab contains the normalized log-base-2 counts of metabolite abundance measured in MuSCs at indicated time post isolation (see methods section for details of data processing).
Cluster1 tab lists the metabolites that are most abundant in freshly isolated MuSCs (see Figure 5C).
Cluster2 tab lists the metabolites that are most abundant in MuSCs 12 hours after isolation (see Figure 5C).
Cluster3 tab lists the metabolites that are most abundant in MuSCs 24 and 48 hours after isolation (see Figure 5C).
BackgroundList tab lists the metabolites that the targeted MS approach is calibrated to detect and serves as the background list for pathway enrichment analyses.
